# Entangled Evolution Model: Application to Adaptive Mutation

**DOI:** 10.1101/2024.09.17.613403

**Authors:** Patrick Ross

**Affiliations:** Independent

## Abstract

Adaptive mutation, the phenomenon where organisms appear to increase beneficial mutation rates in response to environmental stress, has challenged the traditional understanding of random mutations in evolutionary biology. Here, we present a novel computational model, the Entangled Evolution Model, which integrates quantum game theory and quantum walks to provide a new approach to understanding mutation dynamics. By applying the Clauser-Horne-Shimony-Holt (CHSH) quantum game within a modified Lotka-Volterra framework, we simulate how quantum entanglement can accelerate the rate of adaptive mutations in *Escherichia coli* populations. Our simulations reveal that entangled bacterial populations achieve successful adaptive mutations with an 85% probability, significantly surpassing the classical theoretical maximum. These results introduce a new perspective on computational approaches to evolutionary adaptation, offering an efficient algorithmic framework to explore the role of quantum effects in biological systems.

## 1 Introduction

Adaptive mutation challenges the traditional view of mutations as entirely random events in evolutionary biology. Since Cairns et al.’s groundbreaking study “The Origin of Mutants” [1], evidence has accumulated showing that *Escherichia coli* can increase mutation rates in response to environmental stresses, such as nutrient deprivation [2, 3]. These observations suggest that bacteria may possess mechanisms to induce beneficial mutations under specific conditions, prompting a reevaluation of randomness in evolutionary processes.

Recent developments in quantum biology propose that quantum mechanisms, such as quantum entanglement and quantum walks, could influence evolutionary dynamics [4]. Quantum walks (QW) provide a framework for efficient exploration of genotype networks, potentially leading to faster discovery of adaptive mutations compared to classical random walks (CRW). This perspective offers new insights into how quantum coherence and superposition might impact mutation dynamics and adaptation.

In this study, we introduce the “Entangled Evolution” model, integrating the outcomes of the Clauser-Horne-Shimony-Holt (CHSH) quantum game [17] with a modified Lotka–Volterra model. By incorporating principles from quantum game theory and quantum walks, we aim to model adaptive mutation in *E. coli* and demonstrate that quantum entanglement can enhance the probability of beneficial mutations occurring in bacterial populations.

Through these computational simulations, we demonstrate that quantum-enhanced populations can achieve adaptive mutations at a significantly higher rate than classical models would predict. In traditional random mutation models, the probability of a beneficial mutation occurring is extremely small, particularly under environmental stress. However, by incorporating quantum entanglement and using the upper bound from the CHSH quantum game, we show that bacterial populations can effectively communicate and coordinate mutations, achieving the 100-fold increase [2, 3, 1] in mutation rates observed in experimental studies of adaptive mutation. This novel computational approach offers a powerful tool for understanding the impact of quantum effects on evolutionary processes, suggesting that organisms may exploit quantum strategies to optimize their adaptation under stress. The Entangled Evolution Model provides a new pathway for exploring the computational dynamics of evolutionary adaptation and offers testable predictions for experimental studies in quantum biology and evolutionary theory.

## 2 Adaptive Mutation and Its Role in Evolution

Adaptive mutation refers to the observation that bacteria can increase their mutation rates in response to environmental stress, generating beneficial mutations that enhance survival [2, 3]. For instance, in Cairns et al.’s study [1], *E. coli* strains deficient in lactose metabolism produced Lac^+^ revertants at higher frequencies under starvation on lactose media. Subsequent research quantified this increase, showing stressed bacteria exhibit mutation rates up to 100-fold higher than non-stressed cells [10, 11].

Traditional explanations for adaptive mutation involve stress-induced mutagenesis pathways, such as the activation of error-prone DNA polymerases during stress responses [2]. However, these mechanisms do not fully explain the specificity and coordination observed in some experiments, suggesting additional factors may be involved. This opens the possibility for quantum mechanisms, like quantum walks and entanglement, to play a role in facilitating adaptive mutations [4].

## 3 Quantum Mechanics and Evolution

Quantum biology explores the role of quantum phenomena in biological processes. Notably, quantum coherence and tunneling have been implicated in photosynthesis [5, 6], enzyme catalysis [12], and avian magnetoreception [13]. These discoveries challenge the assumption that quantum effects are negligible in warm, noisy biological environments.

In the context of evolution, quantum walks offer a framework for understanding how quantum superposition and entanglement could influence mutation dynamics [4]. Quantum walks are the quantum analog of classical random walks but allow for simultaneous exploration of multiple pathways due to superposition. This property can lead to more efficient searching of genotype networks, potentially increasing the likelihood of discovering beneficial mutations.

Theoretical models suggest that quantum walks can outperform classical random walks in exploring genotype networks, especially in complex, high-dimensional spaces [4]. By applying quantum walks to evolutionary processes, it becomes plausible that organisms could utilize quantum mechanisms to enhance adaptability and survival under stress.

## 4 Entanglement Effects on Bacteria and DNA

The concept of quantum genes posits that DNA nucleotides can exist in a superposition of states due to shared protons in hydrogen bonds, leading to tautomeric shifts that result in mutations [4, 7]. In this framework, the hydrogen bonds between base pairs allow protons to tunnel between nucleotides, creating superpositions that could influence the fidelity of DNA replication.

These quantum superpositions are thought to collapse upon interaction with the environment—a process known as decoherence—resulting in a definitive nucleotide sequence. Environmental factors, such as stress conditions, could affect the decoherence time and bias the outcome toward beneficial mutations [4]. This mechanism provides a potential explanation for the increased mutation rates observed in adaptive mutation experiments.

Maintaining quantum coherence in biological systems is challenging due to thermal fluctuations and interactions with the environment. However, studies have demonstrated that quantum coherence can persist in biological processes at physiological temperatures [5, 6]. These findings suggest that biological systems may have evolved mechanisms to sustain quantum effects long enough to impact biochemical processes, including DNA mutations.

## 5 Utilizing CHSH Quantum Game Theory in Understanding Adaptive Mutation

The CHSH (Clauser-Horne-Shimony-Holt) game serves as a fundamental example in quantum game theory, illustrating how quantum strategies can outperform classical ones in coordination tasks [17, 8]. In the game, two players aim to produce outputs that satisfy a specific condition without communicating, leveraging quantum entanglement to achieve higher success rates.

Drawing an analogy to bacterial populations, each bacterium’s decision to mutate can be likened to a player’s output in the CHSH game. In classical models, mutations occur independently, with a lower probability of simultaneous beneficial mutations across the population. Quantum entanglement could enable bacteria to “coordinate” mutations non-locally, increasing the overall success rate of adaptive mutations [9, 4].

Quantum walks extend this concept by allowing simultaneous exploration of multiple mutational pathways in genotype networks [4]. In contrast to classical random walks, which spread diffusively, quantum walks utilize superposition and interference, leading to faster and more efficient exploration of evolutionary landscapes. This efficiency mirrors the advantage seen in the CHSH game when employing quantum strategies, suggesting that quantum walks could enhance the adaptive capabilities of organisms under stress.

Mathematically, the quantum advantage in the CHSH game is demonstrated by calculating the increased probability of success using entangled states. Similarly, quantum walks on genotype networks show a higher probability of reaching beneficial mutations within a given time frame compared to classical approaches. These models provide a theoretical foundation for the potential role of quantum mechanics in evolutionary biology.

## 6 An Improved Lotka–Volterra Model with Quantum Effects

To model the impact of quantum effects on bacterial population dynamics, we modify the classical Lotka– Volterra competition model. Let *N*_1_(*t*) represent the population of bacteria utilizing quantum-enhanced mutation strategies (entangled bacteria), and *N*_2_(*t*) represent bacteria relying on classical mutation mechanisms (unentangled bacteria). The modified equations are:

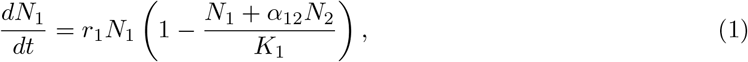

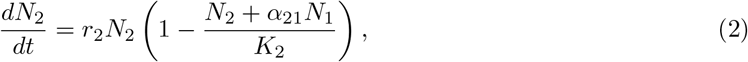

where *r*_*i*_ is the intrinsic growth rate, *K*_*i*_ is the carrying capacity, and *α*_*ij*_ represents the competitive effect of one population on the other.

Incorporating the efficiency of quantum walks in exploring genotype networks, we adjust the competition coefficients to reflect the increased probability of beneficial mutations due to quantum superposition [4]. We set *α*_12_ = 1 − *P*_quantum_ for the entangled population and *α*_21_ = 1 − *P*_classical_ for the unentangled population, where *P*_quantum_ ≈ 0.854 (from the CHSH game and quantum walk simulations) and *P*_classical_ = 0.25 (classical maximum success probability for random mutations under stress).

To reflect the biological scenario where entangled bacteria can utilize lactose while unentangled bacteria cannot, we assign different intrinsic growth rates and carrying capacities:

- *r*_1_ = 0.6, *K*_1_ = 1000 for entangled bacteria.
- *r*_2_ = −0.1, *K*_2_ = 100 for unentangled bacteria.

These values represent the higher growth potential of entangled bacteria due to access to lactose and the decline of unentangled bacteria lacking nutrients.

Simulations using these equations demonstrate that the entangled population outcompetes the unentangled population over time. This outcome aligns with findings from quantum walk models, where the quantum process becomes more efficient at finding beneficial mutations, especially in complex genotype networks [4]. Figure 1 illustrates the population dynamics, highlighting the faster growth and dominance of the entangled bacteria.

**Figure 1.**
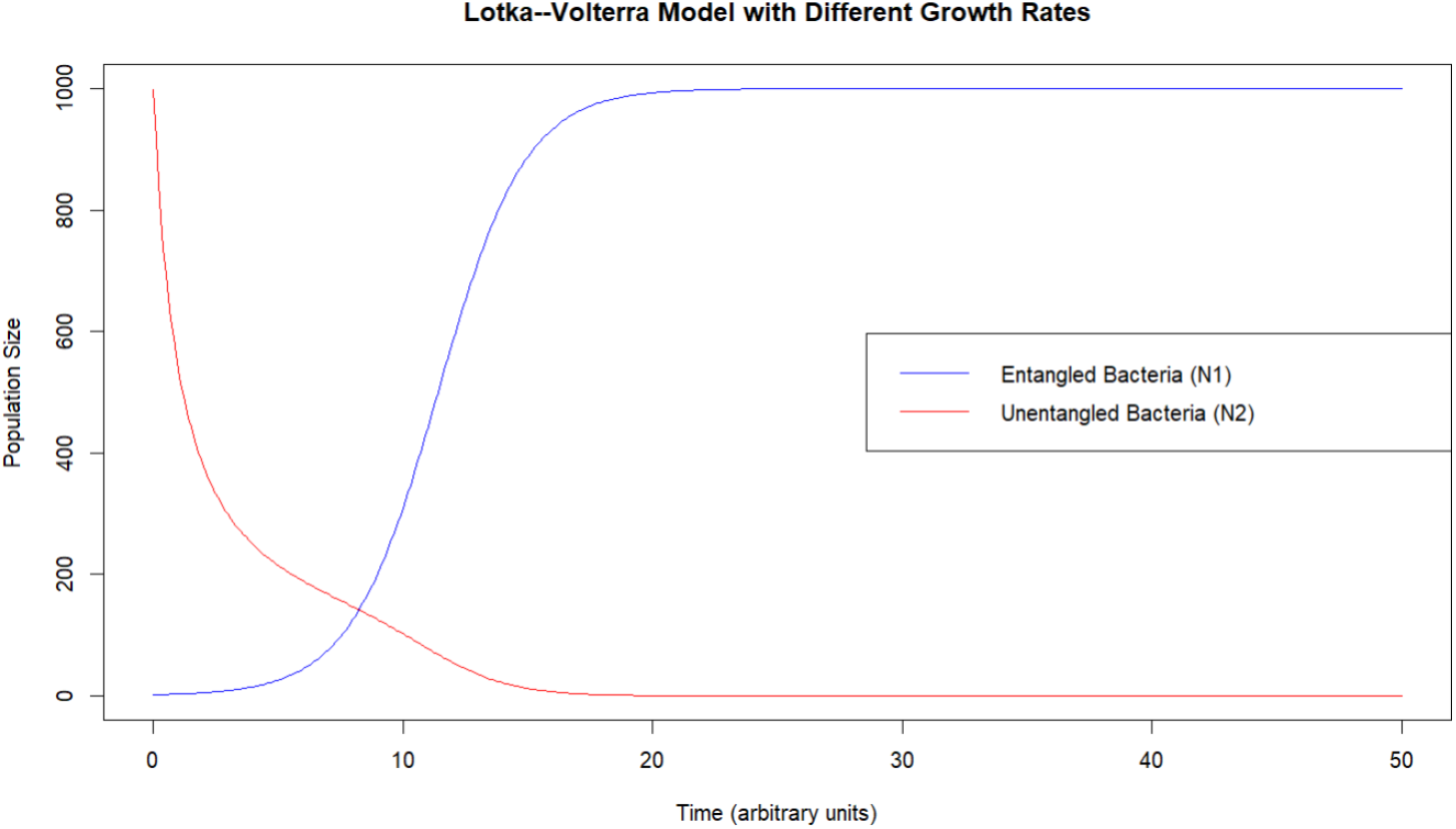
Population dynamics of entangled (*N*_1_) and unentangled (*N*_2_) bacteria modeled by the modified Lotka–Volterra equations. The entangled bacteria rapidly increase in number due to their ability to utilize lactose, while the unentangled bacteria decline over time. See Appendix B for the R script used to generate this figure.

## 7 Discussion

Our findings suggest that integrating quantum mechanics into models of adaptive mutation can account for the increased mutation rates observed in stressed bacterial populations. By incorporating quantum game theory and quantum walks, we provide a theoretical framework where quantum entanglement enhances the likelihood of beneficial mutations, supporting the Entangled Evolution model.

While the application of quantum mechanics to biological systems has faced skepticism, recent theoretical work supports the plausibility of quantum coherence and superposition in biological evolution [4]. Studies have demonstrated that quantum walks can efficiently explore genotype networks, potentially leading to faster adaptation [4]. Moreover, experimental evidence of quantum effects in biological processes, such as photosynthesis and enzyme catalysis, suggests that organisms might exploit quantum mechanisms despite decoherence challenges [5, 6].

To validate our model, we propose experiments using clonal bacterial colonies starting from different positions in the genotype network. By measuring mutation rates and decoherence times under controlled environmental conditions, such as the presence or absence of lactose, we can assess whether quantum mechanisms influence adaptive mutations. Techniques like single-cell sequencing and quantum coherence measurements could provide insights into the role of quantum effects in evolution.

## 8 Philosophical Extensions of Quantum Evolution

The integration of quantum mechanics into evolutionary biology has broader implications for understanding life’s complexity. In phylogenetics, the principle of maximum parsimony suggests that evolution often follows the shortest mutational paths [14]. Quantum walks could provide a mechanism for this efficiency, as they allow simultaneous exploration of multiple evolutionary pathways, potentially explaining the observed parsimony in evolutionary histories [4].

In epigenetics, quantum effects might influence how environmental factors induce heritable changes without altering DNA sequences [15]. Quantum coherence and superposition could play a role in mechanisms like DNA methylation and histone modification, affecting gene expression patterns across generations [4].

Niche construction, where organisms modify their environment, creating new selective pressures [16], could interact with quantum evolutionary processes. Changes in the environment might alter decoherence times or the efficiency of quantum walks in genotype networks, influencing the direction and rate of evolutionary change. This interplay underscores the need to consider quantum effects in ecological and evolutionary studies.

## 9 Conclusion

In conclusion, our Entangled Evolution model bridges quantum mechanics and evolutionary biology, offering a novel explanation for the phenomenon of adaptive mutation. By integrating quantum game theory and quantum walks into population dynamics models, we demonstrate that quantum entanglement could enhance the likelihood of beneficial mutations, providing a potential advantage in survival and adaptation.

This interdisciplinary approach addresses challenges in explaining adaptive mutation solely through classical mechanisms and opens new avenues for research into quantum effects in biological systems. Future work should focus on experimental validation of quantum evolutionary processes and exploring the implications of quantum mechanics in broader biological contexts.

## A Calculations for the CHSH Game

### A.1 Calculations for Maximally Entangled Qubits

To demonstrate the quantum advantage in the CHSH game, we start with the maximally entangled state:

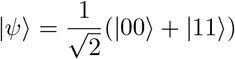

Here, |00⟩ and |11⟩ represent the joint states of the two entangled particles.

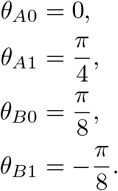

The measurement settings for Alice and Bob are defined by angles *θ*_*A*_ and *θ*_*B*_, corresponding to their inputs *x* and *y*:

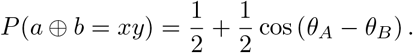

The probability of winning for each combination of inputs is calculated using the quantum correlation functions:

Substituting the measurement angles for each input combination, we find:

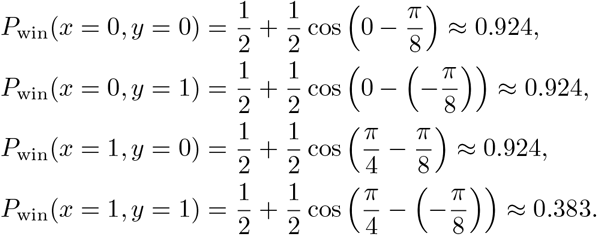

The overall winning probability is then:

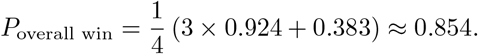

This result shows that by using quantum strategies, Alice and Bob can achieve a success probability of approximately 85.4%, exceeding the classical limit of 75%.

### A.2 Calculations for Sub-Maximally Entangled Qubits

To explore the effects of less-than-maximal entanglement, consider the quantum state coefficients:

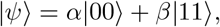

where 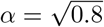 and 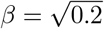

Using the same measurement settings, we calculate the probabilities:

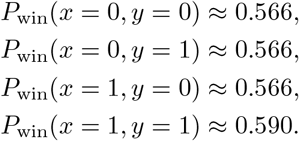

The overall winning probability is:

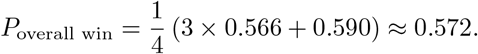

This result indicates that with sub-maximal entanglement, the quantum advantage diminishes, yielding a success probability of approximately 57.2%, which is still above the random classical strategy probability of 50%.

### A.3 Implications for the Model

These calculations demonstrate how the degree of entanglement affects the success probability in the CHSH game. In our modified Lotka–Volterra model, we incorporate these probabilities into the competition coefficients to reflect the enhanced adaptation due to quantum strategies.

For sub-maximally entangled populations:

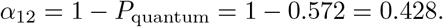

These values are used in Equations (1) and (2) to simulate the population dynamics under different levels of entanglement, illustrating how higher degrees of entanglement confer a greater competitive advantage through more efficient adaptation.

### A.4 Connection to Quantum Walks

The CHSH game calculations align with findings from quantum walk simulations on genotype networks [4]. Quantum walks demonstrate that higher entanglement leads to more efficient exploration of evolutionary pathways, analogous to the increased success probabilities in the CHSH game with greater entanglement.

## B R Script for Lotka–Volterra Simulation with Quantum Effects

The following R script was used to simulate the population dynamics of entangled and unentangled bacteria and generate Figure 1.

**Listing 1:**
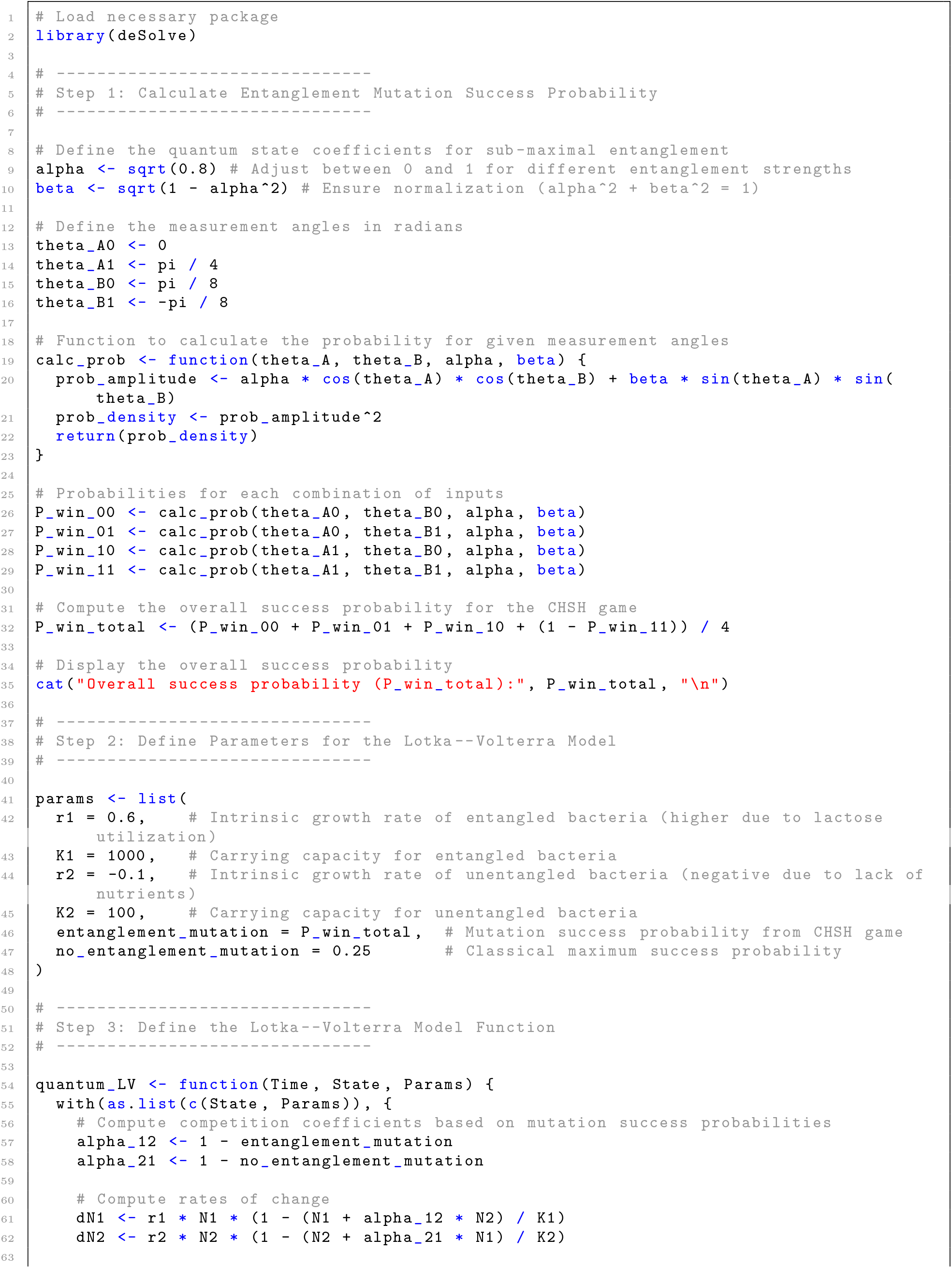
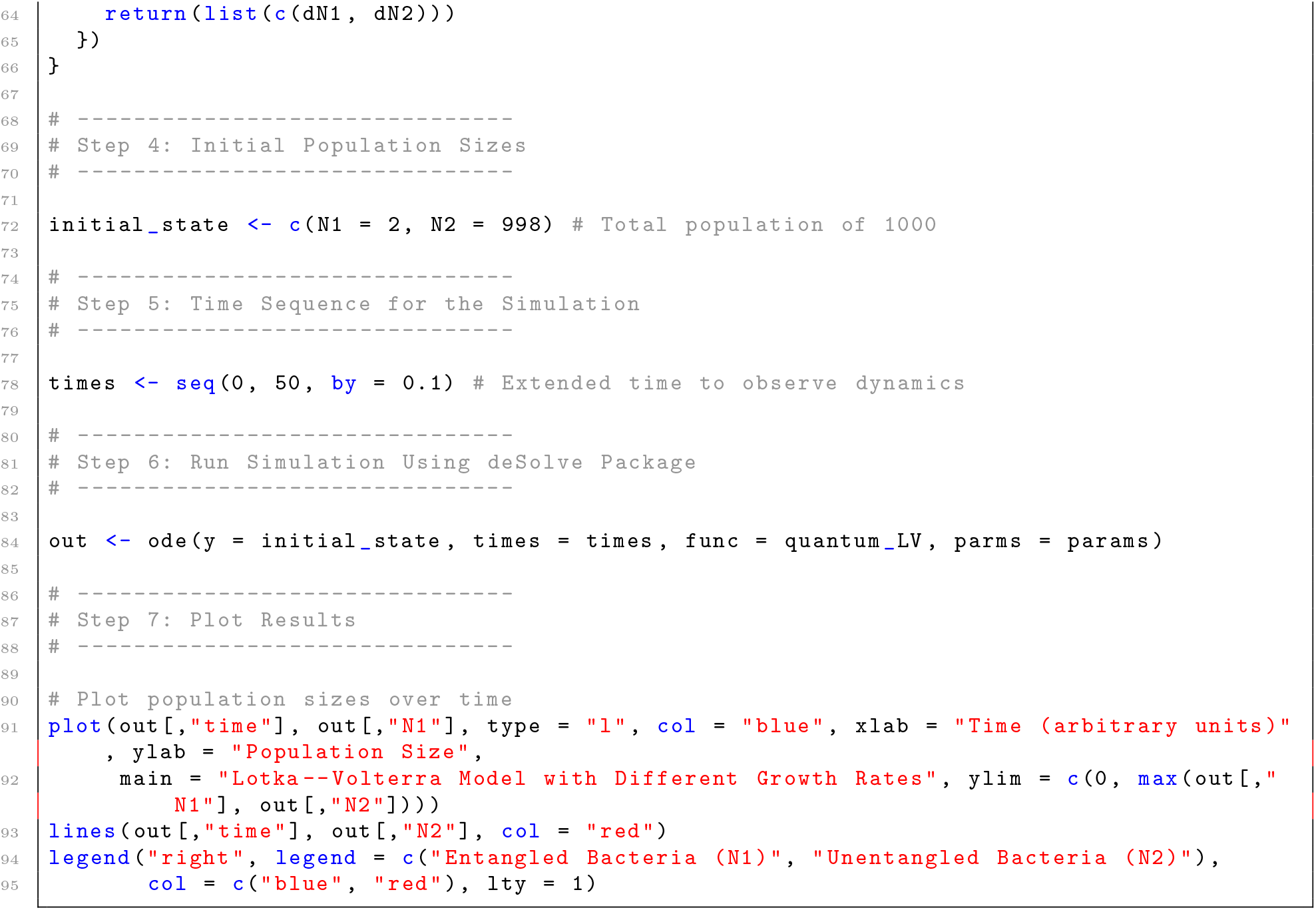
R Script for Lotka–Volterra Simulation with Quantum Effects.

### Notes

- The script calculates the entanglement mutation success probability based on the CHSH game with sub-maximal entanglement.
- Parameters are set to reflect the differing growth rates and carrying capacities of entangled and unentangled bacteria.
- The Lotka–Volterra model function incorporates quantum effects through the competition coefficients.
- The script generates a plot of the population dynamics (Figure 1), demonstrating the advantage of the entangled bacteria.

